# Deep Learning Architectures For the Prediction of YY1-Mediated Chromatin Loops

**DOI:** 10.1101/2022.09.19.508478

**Authors:** Ahtisham Fazeel, Muhammad Nabeel Asim, Johan Trygg, Andreas Dengel, Sheraz Ahmed

## Abstract

YY1-mediated chromatin loops play substantial roles in basic biological processes like gene regulation, cell differentiation, and DNA replication. YY1-mediated chromatin loop prediction is important to understand diverse types of biological processes which may lead to the development of new therapeutics for neurological disorders and cancers. Existing deep learning predictors are capable to predict YY1-mediated chromatin loops in two different cell lines however, they showed limited performance for the prediction of YY1-mediated loops in the same cell lines and suffer significant performance deterioration in cross cell line setting. To provide computational predictors capable of performing large-scale analyses of YY1-mediated loop prediction across multiple cell lines, this paper presents two novel deep learning predictors. The two proposed predictors make use of Word2vec, one hot encoding for sequence representation and long short-term memory, and a convolution neural network along with a gradient flow strategy similar to DenseNet architectures. Both of the predictors are evaluated on two different benchmark datasets of two cell lines HCT116 and K562. Overall the proposed predictors outperform existing DEEPYY1 predictor with an average maximum margin of 4.65%, 7.45% in terms of AUROC, and accuracy, across both of the datases over the independent test sets and 5.1%, 3.2% over 5-fold validation. In terms of cross-cell evaluation, the proposed predictors boast maximum performance enhancements of up to 9.5% and 27.1% in terms of AUROC over HCT116 and K562 datasets.

## I. Introduction

In living organisms proteins are essential to perform diverse types of cellular activities and dysregulation of proteins lead towards development of multifarious diseases such as cancer, neurological, and immunological disorders [1]. Primarily, the production of proteins depends upon the regulation of genes. The process of gene regulation is mediated by different regulatory elements that are distributed in the DNA i.e., enhancers, and promoters. Mainly, extra-cellular signals or transcription factors bind with the enhancer regions to regulate gene expression of nearby or distant genes by forming physically connected chromatin (DNA) loops among enhancers and promoters [2]. These interactions between proximal promoters and distal enhancers often lead to higher order chromatin structure known as topologically associated domains which may contain several chromatin loops [3]. These chromatin loops play a substantial role in performing insulation function to stop the process of transcription.

Different transcription factors are involved in the formation of chromatin loops, such as 11-zinc finger protein (CTCF), and Ying Yang-1 protein (YY1). “Figure 1” illustrates the formation of chromatin loops with the involvement of two different transcription factors CTCF and YY1. YY1-mediated chromatin loops are usually shorter in length, and may bind to smaller motifs of size 12, on the other hand CTCF mediated chromatin loops are larger in length and they form in the regions containing CTCF motifs of 19 nucleotide bases.

**Fig. 1:**
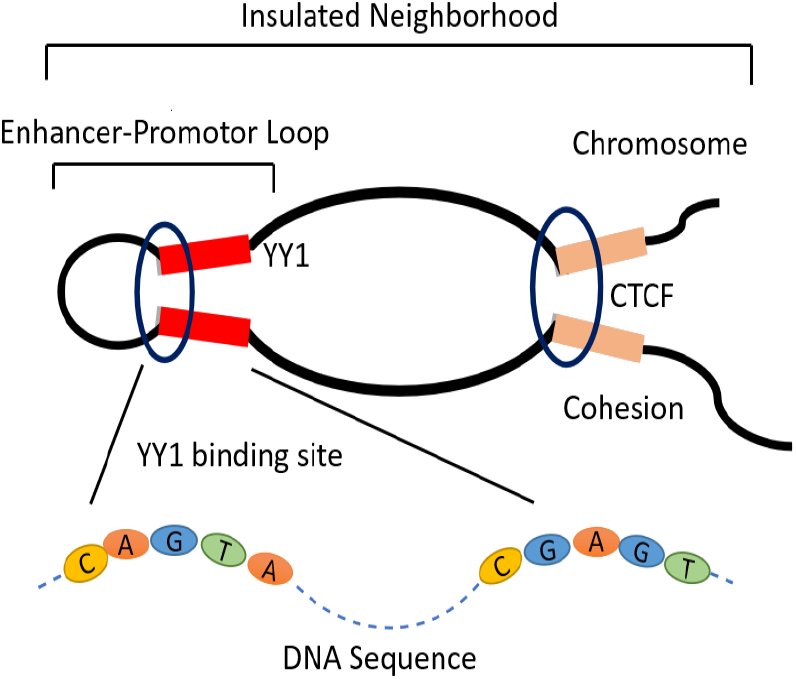
The generic phenomenon of chromatin loop formation due to the interactions between enhancers, promoters, DNA binding proteins (CTCF or YY1), and Cohesion protein.

Several studies reveal the importance of YY1-mediated chromatin loops in certain disorders such as, neurodevelopmental disorders [4], Gabrielede Vries Syndrome [5], ischemic damage, Parkinson and Alzheimer disease [6]. In addition, YY1 can act as a tumor suppressor or stimulator in the case of pancreatic cancer, melanoma, and glioma [6]. These disorders or diseases arise due to the dysregulation at the genetic level caused by the YY1 protein. Despite the importance of gene regulation, cell identity and cell development, the roles of YY1 transcription factor and YY1 mediated chromatin loops are not properly characterized and understood yet.

The identification of generic enhancer promoter interactions provide an abstract level information related to the gene regulation, but such identifications lack information regarding the DNA binding proteins that are involved in initiating such interactions. On the other hand, TF specific enhancer-promoter (EP) interaction and chromatin loops identification can assist in understanding underlying phenomena such as cell-cell communication, and extracellular signalling, which may help out in dealing with complex disorders and cancers or tumors.

Different in-silico and wet lab methods are utilized to identify chromatin interactions and DNA proteins binding sites such as, chromatin interaction analysis by pair-end tagging (ChIA-PET) [7], Chromatin immunoprecipitation (ChIP) [8], High-through chromosome conformation capture (Hi-C), protein-centric chromatin conformation method (HiChIP), chromatin-interacting protein mass spectrometry (ChIP-MS) [6], and proteomics of isolated chromatin segments (PICh) [9]. However, it is laborious, expensive and time consuming to identify chromatin interactions at large number of cell types purely based on such experiments. Particularly, with the avalanche of the sequence data produced at the DNA sequence level, it is highly compelling to develop computational methods for fast, and effective analyses of chromatin loops.

Artificial intelligence (AI) has been a key area of research in genomics to analyze DNA sequence data. Several AI-based methods have been developed with an aim to analyze different chromatin interactions. Majority of these approaches are based on the identification of CTCF mediated loops along with their genomic signatures [10], [11] or generic enhancer promoter interactions. In comparison, there is a scarcity of AI-based predictors for YY1 mediated chromatin loops, for instance only one AI-based method has been developed for the prediction of YY1 mediated chromatin loops namely, DEEPYY1 [12]. DEEPYY1 made use of Word2vec embeddings for encoding DNA sequences and a convolutional neural network for the prediction of YY1 mediated chromatin loops. DEEPYY1 predictor was evaluated on DNA sequence data obtained from HiChip experiments related to two different cell types i.e., HCT116 (colon cell line), and K562 (lymphoblasts from bone marrow). DEEPYY1 predictor managed to produce better performance over same cell line data, however, it failed to produce significant performance over different cell lines data. DEEPYY1 predictor is useful to analyze same cell line based data however, there is a need to develop more generic predictor to perform analysis over different cell lines data.

The paper in hand proposed two deep learning based approaches named densely connected neural network (DCNN) and hybrid neural network (hybrid). Following the observations of different researchers that deep learning based predictors perform better when they are trained on large datasets and considering unavailability of DNA sequences annotated against YY1-mediated chromatin Loops, DCNN-YY1 predictor utilize the idea of transfer learning to generate pretrained k-mer embeddings. Further, in order to extract diverse types of discriminative features, DCNN-YY1 makes use of convolutional layers in two different settings shallow and deep. With an aim to reap the benefits of both types of layers and to avoid gradient vanishing and exploding problems in the process of training, we provide alternative paths for gradient flow among different layers through identity functions which are commonly used in DenseNet architectures [13]. On the other hand, in order to capture positional information of nucleotides the hybrid model makes use of one hot encoding approach for sequence representation. Whereas, the hybrid model itself is comprised of convolution neural network (CNN) and long short terms memory unit (LSTM), to capture discriminative higher spatial and nucleotide level information.

Proposed predictors are evaluated over two different cell lines benchmark datasets. Jointly, over both datasets, experimental results reveals the superiority of the proposed predictors over state-of-the-art DEEPYY1 predictor with average maximum margin of 4.65%, 7.45% in terms of AUROC, and accuracy, across both of the datasets over the independent test sets and 5.1%, 3.2% over 5-fold validation. To explore whether proposed predictors are capable to predict YY1-mediated chromatin loops in different cell lines, we also performed experimentation in cross domain setting in which predictors are trained over sequences of one cell line and evaluation is performed over sequences of other cell line. In cross domain setting proposed predictors outperformed state-of-the-art predictor by a maximum margin of 28% and 10.7% in terms of AUROC, and 22.4%, and 7.01% across accuracy, over HCT116 and K562 datasets.

## II. Material and Methods

This section briefly demonstrates the working paradigm of the proposed YY1 chromatin loop predictors, benchmark datasets, and the evaluation metrics.

### A. CNN and LSTM Based YY1-Mediated Chromatin Loop Predictor

The working paradigm of the proposed hybrid (CNN+LSTM) model can be divided in two main stages. At the first stage, one hot encoded vectors are generated from the nucleotides of DNA sequences. At the second stage, the hybrid model utilizes these vectors to predict YY1-mediated chromatin loops. The working paradigm of one hot encoding and the hybrid model are discussed in the following sections.

#### 1) One Hot Encoding

One hot encoding is a simplified yet effective way of representing genomic sequences for classification, which may encode the nucleotide composition information. In this method, out of four different nucleotides, each nucleotide is represented by a vector of size 4 as shown below,

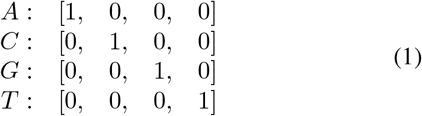

**Fig. 2:**
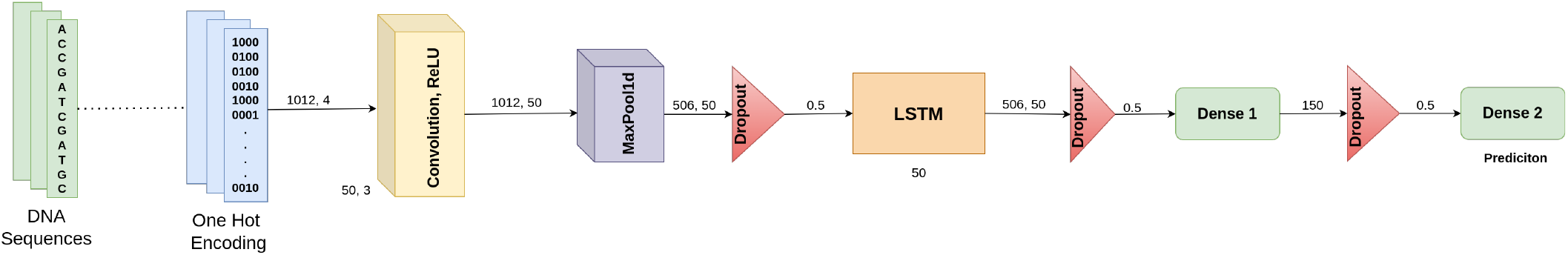
The graphical illustration of the proposed (hybrid) architecture.

#### 2) LSTM and CNN

Convolutional neural networks (CNNs) are used widely in the domain of computer vision and natural language processing [14]. A CNN is comprised of three different layers: convolution, pooling, and fully connected layers, which allow it to capture spatial features from the input. The convolution operation leads to the formation of feature maps, whereas the pooling operation reduces the dimension of the feature maps by taking either average or maximum value. CNNs are commonly used for DNA analysis problems to learn local features such as, motifs in the case of DNA sequences and are often referred as local feature learning layers. On the other hand, CNN ignores the dependence present within the inputs which deteriorates their power to model NLP-based problems accurately.

Therefore, recurrent neural networks and their variants i.e., long short-term memory (LSTM), and gated recurring units (GRUs) are used to learn such long-term dependencies. LSTM contains a series of memory cells, which are dependent on three different gates to compute the output i.e., input, forget, and output gate. The input gate adds the input to the current cell state by the use of two non-linear activation functions i.e., sigmoid and tanh. Sigmoid assigns a probability to the inputs where tanh transforms them in the range of −1 and 1. The forget gate is responsible for the removal of undesired information from the inputs, it achieves this by taking two different inputs i.e., *h*_*t*–1_, and *X_t_*. These inputs are multiplied with the weights and a bias is added in them. The output of the multiplication operation is followed by a sigmoid operation that transforms these values in the range of 0 (forget) and 1 (remember). The output gate gives the output of the LSTM cell based on the sigmoid and tanh activation functions.

In the current setting, we make use of one hot encoding to represent DNA sequences in the form of vectors. One hot encoded DNA sequences are passed through the convolution, and max-pooling layers to extract the local features, which is followed by the LSTM layer to learn long-term sequence dependencies and a fully connected layer for classification.

### B. Densely Connected Neural Network Based YY1-Mediated Chromatin Loop Predictor

The working paradigm of the proposed DCNN-YY1 predictor can be divided in two different stages. At the first stage, pretrained k-mer embeddings are generated in an unsupervised manner using well known Word2vec model [15]. In second stage, DCNN-YY1 utilizes pretrained k-mer embeddings and raw DNA sequence to predict YY1-mediated chromatin loops.

The working paradigm of Word2vec algorithm is summarized in the subsection II-B1. Furthermore, CNNs and dense connectivity are comprehensively discussed in subsection II-B2.

#### 1) Word2vec

Word2vec is a two-layered neural network model that is capable to learn associations of k-mers from the raw DNA sequence data [16]. Word2vec takes DNA sequences as an input and transforms the sequence data into a numerical vector space, where each k-mer is represented by a N-dimensional vector. Such vectors include important characteristics related to k-mers with respect to four unique nucleotides i.e., semantic relationships and contexts. Moreover, the k-mers that are similar or semantically close to each other they lie closer in the continuous vector space.

Two common methods are used in a Word2vec model for the generation of embeddings namely, common bag of words (CBOW), and skipgram [16]. CBOW works by predicting the target k-mer when provided with the distributed representations of the sequence k-mers. Whereas, the skipgram model tries to predict the context of a k-mer which is opposite to the working paradigm of CBOW model.

We generate 7 different overlapping k-mers from range 1 to 7. Iteratively, for each k-mer we generate 100 dimensional k-mer embeddings using CBOW model. Based on the size of k-mer we obtain 100 dimensional vectors associated to each unique k-mer. For instance, for 1-mer there exist 4 unique 1-mers A, C, G, T, so we obtain 4, 100 dimensional vectors. For 2-mers we obtain 16 and for 3-mers we obtain 64 vectors and so on. K-mer embeddings are generated separately for enhancer and promoter sequences.

#### 2) Convolutional Neural Networks and Dense Connectivity

We utilize CNN based architecture for YY1-mediated chromatin loop prediction, “Figure 3” shows the complete architecture of the proposed predictor. The network consists of three 1-dimensional (1-D) convolutions, 2 dropout and 4 fully connected layers.

**Fig. 3:**
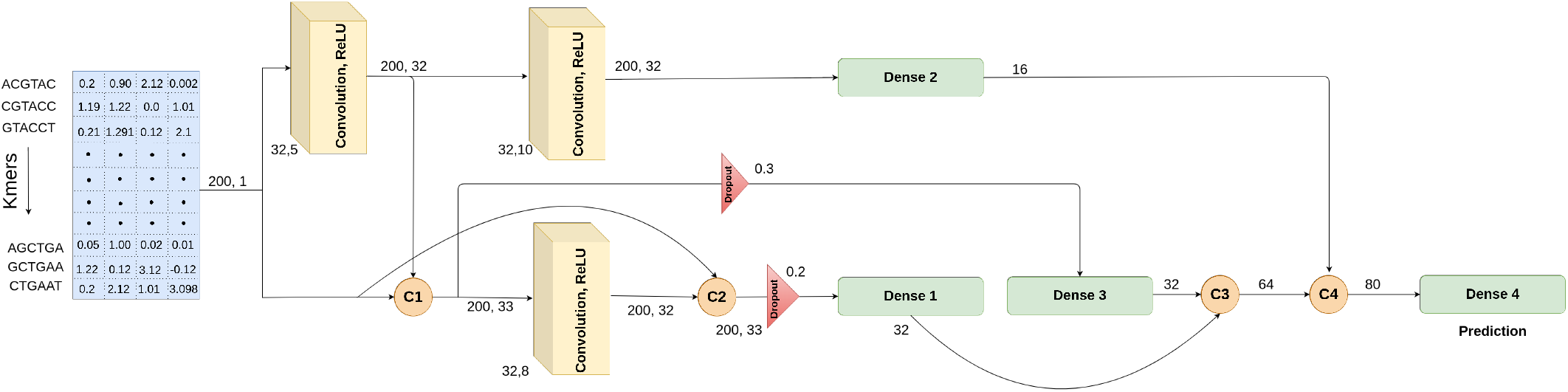
The graphical illustration of the proposed (DCNN) architecture.

We generate k-mers of enhancer and promoter sequences and transform enhancer k-mer sequences to 100 dimensional statistical vectors by taking average of k-mers pretrained vectors that are generated over enhancer sequences. Similarly, promoter k-mer sequences are transformed to 100 dimensional vectors by utilizing precomputed k-mer vectors over promoter sequences. Statistical vectors of both sequences are concatenated to generate single 200 dimensional vector for each sample. This statistical vector is passed through 1-D convolutional layers to extract robust and meaningful features for further processing i.e, the output size (200 × 32) remains same across each convolution due to the usage of paddings. Each 1-D convolution layer is followed by an activation layer which uses rectified linear unit (ReLU) as an activation function throughout the network except for the last fully connected layer which utilizes Sigmoid function for binary classification.

We further amplify the representational power of the proposed YY1-mediated chromatin loop predictor by the use of dense connectivity that is inspired by the concept of identity mapping or skip connections [17]. Skips connections allow to train the network in a more efficient way. In skip connections the input of a layer is added to the output of a layer, but in terms of dense connections [13] the input of a layer is concatenated to the output of the layer thus offering multiple advantages such as, less vanishing-gradient problem, better feature propagation, and substantial reduction in the number of parameters. “Figure 3” illustrates 4 dense connections namely, C1, C2, C3, and C4 for better feature propagation throughout the network.

### C. Experimental Setup

The proposed predictors are implemented in Keras. Adam is used as an optimizer with a learning rate of 0.01, and binary cross-entropy is used as the loss function. For the experiments of this study, the DCNN model is trained only for 6 epochs with a batch size of 32. Whereas, the hybrid model is trained for 20 epochs in cross-validation and independent test settings. In addition, the parameters of CNN and LSTM layers are provided in the Figures 3, and 2.

### D. Dataset

We utilized the datasets of DEEPYY1 [12] related to two different cell lines i.e., K562, and HCT116. The datasets were collected from the HiChip experiments of Weintrub et al., [18]. As the details related to the preprocessing of the DNA sequences are given in the study of DEEPYY1 [12], therefore we summarize here the number of positive and negative samples in the train and test sets of HCT116 and K562 cell lines. The datasets of both cells are well balanced in terms of positive and negative samples, and “Table I” demonstrates the number of positive and negative samples in train and test sets of HCT116 and K562.

**TABLE I:** Statistics of benchmark datasets.

| Cell Type | Set | +EPIs | -EPIs |
| --- | --- | --- | --- |
| K562 | Train | 3863 | 3866 |
|  | Test | 1657 | 1657 |
| HCT116 | Train | 2095 | 2097 |
|  | Test | 898 | 898 |

## III. Evaluation

To evaluate the integrity and predictive performance of the proposed predictor, following the evaluation criteria of the state-of-the-art [12], we utilized two different measures, i.e., accuracy (ACC), and area under the receiver operating characteristic curve (AUROC).

Accuracy (ACC) measures the proportion of correct predictions in relation to all predictions. Area under receiver operating characteristics (AUROC) calculates performance score by using true positive rate (TPR) and false positive rate (FPR) at different thresholds, where true positive rate (TPR) gives the proportion of correct predictions in predictions of positive class and false positive rate (FPR) is the proportion of false positives among all positive predictions (the sum of false positives and true negatives).

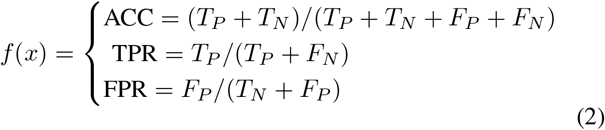

In the above cases, *T_P_* denotes the true positive predictions, *T_N_* shows true negative predictions. Whereas, *F_P_* and *F_N_* refer to the false predictions related to the positive and negative classes.

## IV. Results

This section demonstrates the performance of the proposed and state-of-the-art DEEPYY1 [13] predictor using two benchmark datasets sets in 5-fold cross-validation setting and independent test sets.

### A. Intrinsic Analyses of DNA Sequence Representations

To perform an intrinsic analysis of the sequence representations obtained by the word2vec-based model, we chose both the benchmark datasets and plot their feature matrix using the t-distributed stochastic neighbor embedding (TSNE) method. Figure 4 shows the embedding space for the samples belonging to both classes (chromatin loop and non-chromatin loop). In addition, the representation of positive and negative class samples leads to the formation of unique yet slight dependent clusters, which suggests that the sequence representations are discriminatory enough to be used for the purpose of the YY1-mediated chromatin loop prediction across HCT116 and K562 datasets.

**Fig. 4:**
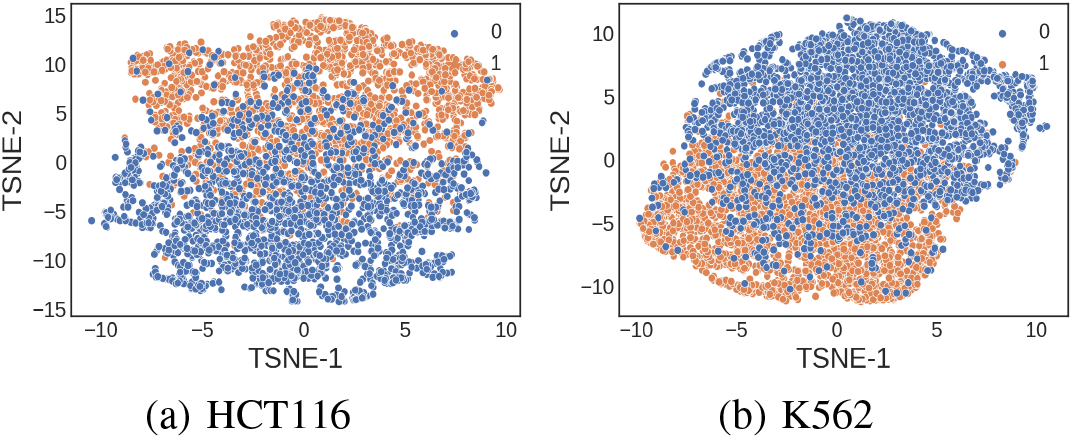
TSNE visualization of the word2vec based embeddings used to train DCNN architecture for YY1-mediated chromatin loop prediction.

### B. Proposed DCNN Predictor Performance

Figure 5 shows the effect of different k-mers on the performance values of the proposed predictor. Lower-sized k-mers yield frequent and less unique patterns in the DNA sequences which lead to low performance of the proposed predictor i.e., *K* = 1, ···, 3. As, K-mer size is increased the performance scores also increase, where the performance scores are maximum at K=6, as higher k-mers lead to unique patterns in the DNA sequences which are crucial for the generation of discriminative sequence representations. For K=7 the performance deteriorates due to the formation of rare patterns. Hence, the value of K=6 is selected for further experiments and performance comparisons.

**Fig. 5:**
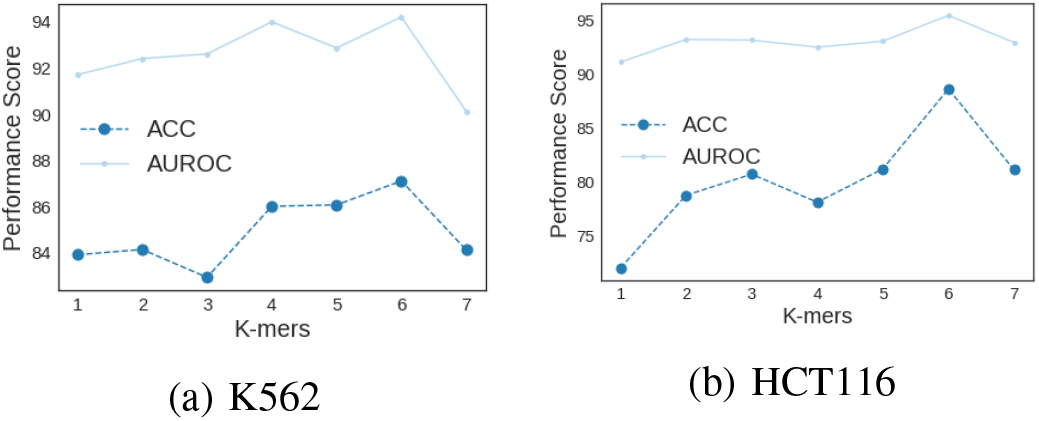
Proposed predictor performance analyses at different k-mers.

### C. Predictive Performance Analyses Over K562 and HCT116

Figures 6 and 7 compare accuracy and AUROC values produced by proposed (hybrid, and DCNN) and state-of-the-art DEEPYY1 predictors at 5-fold cross-validation and independent test sets

**Fig. 6:**
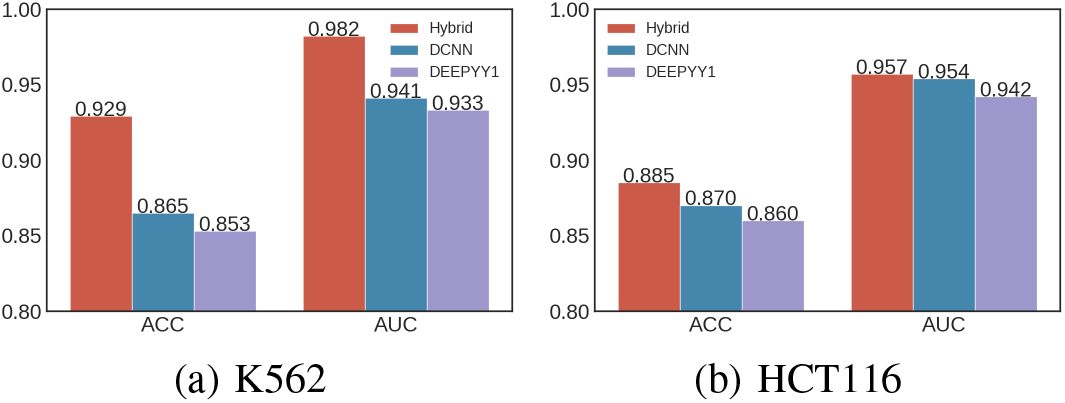
Performance comparison of the proposed and state-of-the-art predictors on two benchmark datasets using 5-fold cross-validation.

**Fig. 7:**
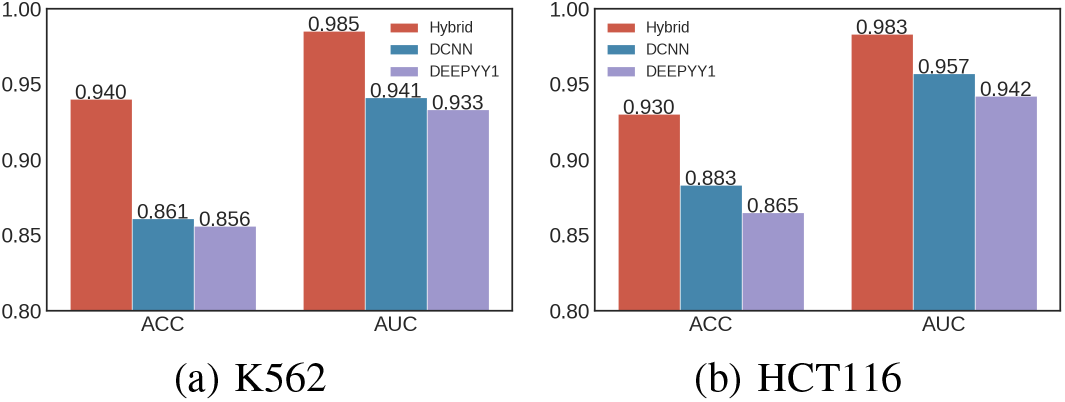
Performance comparison of the proposed and state-of-the-art predictors on two independent test sets.

Figure 6 shows that the proposed hybrid (CNN+LSTM) predictor achieves AUROC values of 98.2% and 95.7% across K562 and HCT116 datasets. Whereas, in terms of accuracy the proposed predictor achieves 92.9% and 88.5% across K562 and HCT116 datasets. Overall, in comparison to the state-of-the-art DEEPYY1 [12], the proposed predictor shows performance improvements in terms of AUROC and ACC with the average margins of 3.02% and 5.05% across both K5652 and HCT116 datasets, via 5-fold cross-validation. Figure 6 shows that the proposed DCNN predictor achieves AUROC values of 94.1% and 95.4% across K562 and HCT116. Whereas, in terms of accuracy the proposed predictor achieves 86.5% and 87.0% across K562 and HCT116 datasets. Overall, in comparison to the state-of-the-art DEEPYY1 [12], the proposed predictor shows performance improvements across AUROC and ACC with the average margin of 1% and 1.1% across both of the datasets, via 5-fold cross-validation.

Figure 7 shows proposed hybrid (CNN+LSTM) predictor produces 98.5%, and 98.3% AUROC values over K562 and HCT116 independent test sets. Whereas, in terms of accuracy the proposed hybrid predictor shows ACC values of 94.0% and 93.0%. Overall, the proposed predictor provides performance improvements in terms of ACC and AUROC with significant margins i.e., 7.45% and 4.65% in comparison to the state-of-the-art DEEPYY1 predictor. Figure 7 shows proposed DCNN predictor produced 94.1%, and 95.7% AUROC values over K562 and HCT116 independent test sets. Whereas, in terms of accuracy the proposed DCNN predictor shows ACC values of 86.1% and 88.3%. Overall, the proposed predictor provides average performance improvements over ACC and AUROC i.e., 2.3% and 1.15% in comparison to the state-of-the-art DEEPYY1 predictor.

### D. Performance Analyses Over Cross Cell Data

To assess the generalizability of the models over different cell lines data, we train the models for chromatin loops in a cross-cell manner such that the models are trained on K562 cell data and predictions are performed on second cell HCT116 data and vice versa. Table II demonstrates the AUROC and ACC performance values of proposed and DEEPYY1 predictors.

**TABLE II:** Performance values of the proposed and existing predictors on the basis cross cell line testing.

| Method | Training Data | Testing Data | Accuracy | AUC |
| --- | --- | --- | --- | --- |
| **DEEP-YY1** | K562 | HCT116 | 80.0 | 87.9 |
|  | HCT116 | K562 | 66.0 | 70.1 |
| **DCNN** | K562 | HCT116 | 86.0 | 94.1 |
|  | HCT116 | K562 | 84.2 | 92.5 |
| **Hybrid** | K562 | HCT116 | **91.0** | **97.4** |
|  | HCT116 | K562 | **91.2** | **97.2** |

The proposed hybrid predictor outperforms DEEPYY1 by a margin of 9.5% in terms of AUROC over train-test set of K562-HCT116 and 27.1% over a train-test set of HCT116-K562. Overall, the proposed hybrid predictor shows consistent and better performance in cross-cell evaluation in terms of AUROC. Similarly, in terms of accuracy, the proposed hybrid predictor outperforms DEEPYY1 by 11.0% over HCT116 test set and 25.2% over K562 test set, which shows the generalizability power of the proposed approach.

The proposed DCNN predictor outperforms DEEPYY1 by a margin of 7.01% in terms of AUROC over train-test set of K562-HCT116 and 22.4% over train-test set of HCT116-K562. Overall, the proposed DCNN predictor shows consistent and better performance in cross cell evaluation in terms of AUROC. Similarly, in terms of accuracy, the proposed predictor outperforms DEEPYY1 by 6% over HCT116 test set and 23.2% over K562 test set, which shows the generalizability power of the proposed approach.

The better performance offered by the proposed DCNN predictor is subjected to the use of dense connectivity in the CNN, as it allows the model to learn in a more suitable manner due to the better feature propagation in the deeper layers. In comparison, the lower performance of the DEEPYY1 is because of the 1 layer CNN which does not possess enough learning power for complex genomic sequences, and the use of max-pool layer which often ignores very crucial information in terms of textual data. Similarly, the proposed hybrid approach takes in to account nucleotide composition information and learns the dependencies of nucleotides with an incorporated LSTM which makes it superior to the other approaches.

## V. Conclusion

This study presents two new YY1-mediated chromatin loop predictors based on CNNs, and RNNs and dense connectivity. It illustrates the impact of different k-mers on the predictive performance of the proposed DCNN predictor, where 6-mer lead to the best performance. The analyses shows that both the proposed predictors are able to generalize well on similar and cross-cell datasets. It also demonstrates that the proposed predictors offer performance superiority over the state-of-the-art DEEPYY1. Overall the proposed predictors outperform existing DEEPYY1 predictor with an average maximum margin of 4.65%, 7.45% in terms of AUROC, and accuracy, across both of the datases over the independent test sets and 5.1%, 3.2% over 5-fold validation. In terms of cross-cell evaluation, the proposed predictors boast maximum performance enhancements of up to 9.5% and 27.1% in terms of AUROC over HCT116 and K562 datasets. The proposed predictors can assist in understanding the process of transcriptional regulation, and multiple disorders which are related the YY1-mediated chromatin loops. Furthermore, in the future, this approach can be leveraged for large-scale cellular chromatin loops analyses and also for other chromatin loops predictions.

